# Quod erat demonstrandum? No restriction endonuclease fold in MIF

**DOI:** 10.1101/085258

**Authors:** Lakshminarayan M. Iyer, L. Aravind

## Abstract

It was claimed in a recently published article that MIF functions as an exo/endo-DNase mediating cell-death upon being induced by DNA damage and PARP1. MIF, for which tautomerase activity has been previously reported, is a member of the tautomerase superfamily which does not feature nucleases. The central premise of the authors to suggest that MIF functions as a DNase is the supposed structural relationship to nucleases of the Restriction endonuclease (REase) fold, which frequently but not always contain a motif of the form PD-(D/E)XK. However, we present evidence to show that this claim is entirely unsupported.

---

In a recent publication it was claimed that the human protein MIF functions as an exo/endo-DNase mediating cell-death upon being induced by DNA damage and PARP1 (1). MIF is a member of the tautomerase superfamily which does not feature nucleases (2, 3). In line with this, previous studies have reported tautomerase activity for human MIF (2, 3). The central premise of the authors to suggest that MIF functions as a DNase is the supposed structural relationship to nucleases of the Restriction endonuclease (REase) fold (1), which frequently but not always contain a motif of the form PD-(D/E)XK (4). Further, they claim that MIF contains three copies of this motif implying that it contains three copies of the REase fold domain. However, a reanalysis of the MIF sequence suggests that this claim is not borne out by any kind of structure or sequence evidence. Additionally, there are several elements of structural and functional misunderstanding in the original claim. Accordingly we present below a brief analysis to indicate why this is so.

The following lines of structural evidence strongly disfavor MIF having any relationship to the REase fold enzymes: 1) A DALI search with the structure of MIF does not recover any REase fold structures with Z-scores suggestive of genuine relationships(Z>3) (5). However, as expected it recovers several tautomerase superfamily structures (5) which reaffirms its originally recognized relationship. 2) The REase fold is well-characterized and always contains a core sheet of five strands and two □-helices which pack against opposite sides of the sheet (Fig. 1A) (4), (5). As it lacks internal symmetry there is no structural evidence that it arose from the duplication of a simpler element. In contrast, the tautomerase fold (Fig. 1A, B) has clear internal symmetry, comprising of two copies of a simple structural element with two core strands and an □-helix. This unit also occurs as a standalone and the catalytic residues are in several cases symmetrically distributed in both the units (3, 6). Thus,there is no doubt that in contrast to the REase fold, the tautomerase fold has emerged from an internal duplication of a simple unit making it topologically unrelated to the former fold (4). 3) The REase fold catalysis requires additional residues (not conserved in MIF) beyond the two metal-coordinating acidic residues of the so-called PD-(D/E)XK motif (6). Moreover, the aspartates and glutamates identified by the authors (1) are often on opposite ends of different structural elements of the tautomerase fold and not proximal in space to coordinate a metal ion (Fig. 1A) (5). 4) Profile-profile searches (7) with hidden Markov models based on both diverse REase fold members and tautomerase superfamily protein never recover each other. 5) Finally, several residues in the so-called triplet of PD-(D/E)XK motifs identified by the authors (1), unlike the characteristic prolines and other catalytic residues of the tautomerases (3), are not entirely conserved even among animal orthologs, much less those from other eukaryotes and prokaryotes (5) (Fig. 1C). This indicates a selective pressure primarily acting to preserve the tautomerase activity.

**Fig. 1.**
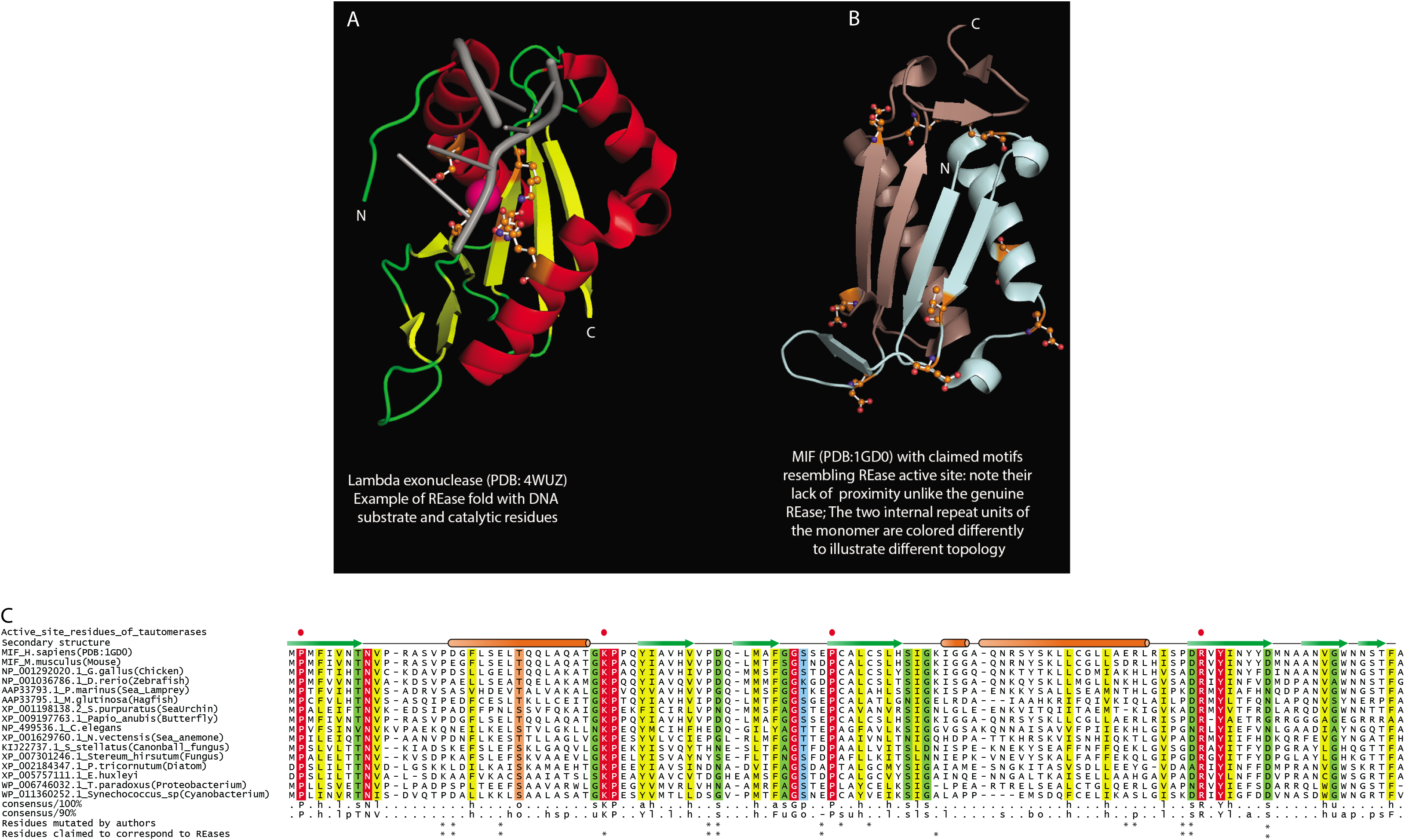
Structures and sequences of the restriction endonuclease fold and MI. (A) Structure of a classic representative of the REase fold (the phage lambda exonuclease). (B) Structure of the MIF tautomerase. (C) Multiple sequence alignment of representative orthologs of MIF showing the conserved tautomerase active site and the other less-conserved residues claimed to form the DNase active site.

Further structural mischaracterization of MIF arises from the authors' claim linking glutamate 22 to the catalytic residue of the so-called Exonuclease-Endonuclease-Phosphatase (synaptojanin-like) domain (Pfam: PF03372). This residue is seen by the authors as being equivalent to the glutamate in the PD-(D/E)XK motif. However, the synaptojanin-like fold is unrelated to both the REases with the so-called PD-(D/E)XK motif as well as the tautomerases (4, 8). Hence, making an argument for the catalytic role of this residue in MIF based on the synaptojanin-like domain arises from the conflation of residues from unrelated folds and is entirely unwarranted. This also appears inconsistent with the experimental results as the authors mutate this glutamate as primary evidence for their claim. They accordingly state that mutating it to alanine has less deleterious effects than mutating it to glutamine. In genuine REase fold enzymes the E->Q substitution is widely seen with active enzymes (4) whereas A in the same position results in an inactive enzyme. Similar mutating the conserved glutamate to an alanine in the synaptojanin-like domain should have more deleterious effects.

Moreover, the authors claim that MIF contains a "CxxCxxHx(n)C Zinc finger" which they claim to
be found in DNA repair proteins like the Fanconi anemia protein FAN1/KIAA1018. The supposed
Zn-chelating motif occurs at residues 60 to 84 in the sequence of MIF (Fig. 1C). First, the sequence in MIF is unrelated to the Rad18 Zn-finger present in FAN1/KIAA1018 as the Zn-chelating histidine in that protein is not positioned 2 residues after the second cysteine. Consistent with this they do not recover each other in any type of sequence profile search (9). Indeed, none of the metal-chelating residues of the sequence presented by the authors are strongly conserved even among MIF orthologs indicating that there is no selection for any metal-coordination function. More tellingly, an examination of the structure of MIF shows that the side chains of these residues are nowhere proximal to effectively coordinate a Zn ion. Thus, the use of this sequence as an argument for a DNA-related function is flawed.

Hence, this demonstration of DNase function in MIF is based on a flawed structural hypothesis;
accordingly, its metal-dependent DNase activity should be viewed as being potentially suspect.

